# Lesion network localization of a stable personality trait

**DOI:** 10.1101/2023.06.05.543261

**Authors:** Stephan T. Palm, Sanaz Khosravani, Fabio Campanella, Alexander L. Cohen, William Drew, Franco Fabbro, Roger Pottanat, Cosimo Urgesi, Michael A. Ferguson, Shan H. Siddiqi

## Abstract

**Objective:** To identify a convergent neuroanatomical substrate for novelty seeking (NS), a stable personality trait, which could serve as a therapeutic target for transcranial magnetic stimulation (TMS).

**Methods:** We analyzed two independent datasets (total n=136) in which the Temperament and Character Inventory (TCI) was used to document alterations across 7 stable personality traits before and after brain lesions surgically induced by glioma resection. Using lesion network mapping, we examined whether alterations in NS were causally associated with lesions to specific brain networks. We assessed for strength of replication across datasets and specificity against other personality traits.

**Results:** Lesion locations that are linked to NS map to a common brain with its largest cluster in the dorsal cingulate. This map was specific to NS relative to other personality traits and overlapped with prior published neuroimaging findings related to the term “novelty”. Utilizing a pre-computed connectome, we also derived a map highlighting potential targets for non-invasive brain stimulation that may alter this stable personality trait.

**Conclusions:** We derived and cross-validated a brain network that is functionally connected to lesions that are causally responsible for the stable personality trait “novelty seeking”. Lesions to this network were associated with changes in NS. This includes a superficial node within the dorsolateral prefrontal cortex that may serve as a promising TMS target to modulate or protect against abnormal NS.

## Introduction

The neurobiological basis of personality traits has been a subject of ongoing debate. While earlier theories predominantly emphasized the role of environmental factors in personality development, recent research has provided compelling evidence elucidating the fundamental role of genetics and neuroanatomy (Gartstein et al., 2016). One existing construct for investigating the neurobiological foundations of personality is the Temperament and Character Inventory (TCI). Diverging from personality measures that are primarily derived from population-based clustering, the TCI adopts a biopsychosocial framework that considers the intricate interactions among genetic, psychological, social, cultural, and spiritual aspects (Cloninger & Cloninger, 2011; Cloninger & Garcia, 2015). This includes four temperament traits that are associated with heritable neurobiology, as well as three character traits that are related to social learning.

Novelty seeking (NS) is a temperament trait referring to intensity of exhilaration or excitement in response to novel stimuli (Cloninger et al., 1991). High NS predicts risky behaviors such as substance use disorders (Bardo et al., 1996; Grucza et al., 2006; Stansfield & Kirstein, 2006), eating disorders (Krug et al., 2009), and impulsive behaviors (Arenas & Manzanedo, 2017). Patients with borderline personality disorder tend to have exaggerated novelty seeking, while patients with obsessive-compulsive personality disorder tend to have impaired novelty seeking (*Personality and Psychopathology*, 1999). Consequently, investigation of the neural correlates of NS has been a focus of cognitive neuroscience and psychiatric research. Correlative neuroimaging studies have associated novelty seeking with the left middle frontal gyrus, posterior cingulate cortex, and ventral striatum (Del Giacco et al., 2022; Silverman et al., 2015). However, it remains unclear if there is a brain network that is causally implicated in novelty seeking.

More recently, focal brain stimulation has emerged as a promising avenue for modifying the underlying neural mechanisms of neuropsychiatric disorders. These techniques involve selectively activating or inhibiting specific brain networks that are causally implicated in the underlying symptoms or disease processes. This is most commonly applied for major depression but has also been used to modulate stable personality traits (Faerman et al., 2021). However, the effects of neuromodulation are target-dependent and the optimal targets for NS remain unclear. Recent advancements in causal brain mapping techniques (Fox, 2018) have shown that spatially diverse lesions often share common connectivity, offering novel insights into the neuroanatomy of these lesion-induced syndromes (Siddiqi et al., 2022). Across multiple different symptoms and syndromes, lesions causing a particular symptom tend to share a common connectivity profile with stimulation sites that relieve the same symptom (Ganos et al., 2022; Schaper et al., 2023; Siddiqi et al., 2021). To extend this approach and identify brain stimulation targets for a stable personality trait, we mapped the connectivity of brain lesions that modify novelty seeking across two datasets. We hypothesized that a consistent and specific network would be identified.

## Results

### Lesions modifying novelty seeking are connected to a common brain network

We identified two independent datasets (n=88 and n=48) in which the TCI was collected before and after a glioma resection (Campanella et al., 2014; Urgesi et al., 2010). Lesions to different brain regions led to differential increases or decreases in NS (Fig. 1). To map the network associated with lesion-induced changes in novelty seeking, we first mapped each lesion (Fig. 2a) to its underlying connectivity profile (Fig. 2b) using a normative connectome database generated from 1000 healthy controls, as illustrated in our prior work (A. Cohen et al., 2021; A. L. Cohen & Fox, 2020; Fox, 2018; Siddiqi et al., 2021). The resultant connectivity profiles (t-maps) were subsequently compared with changes in the subscales of TCI across all study participants using Pearson correlations at each voxel (Fig. 2c). This yielded distinctive brain network maps depicting the connectivity of brain lesions that are more likely to modify novelty seeking (Fig. 2d) and other traits.

**Figure 1:**
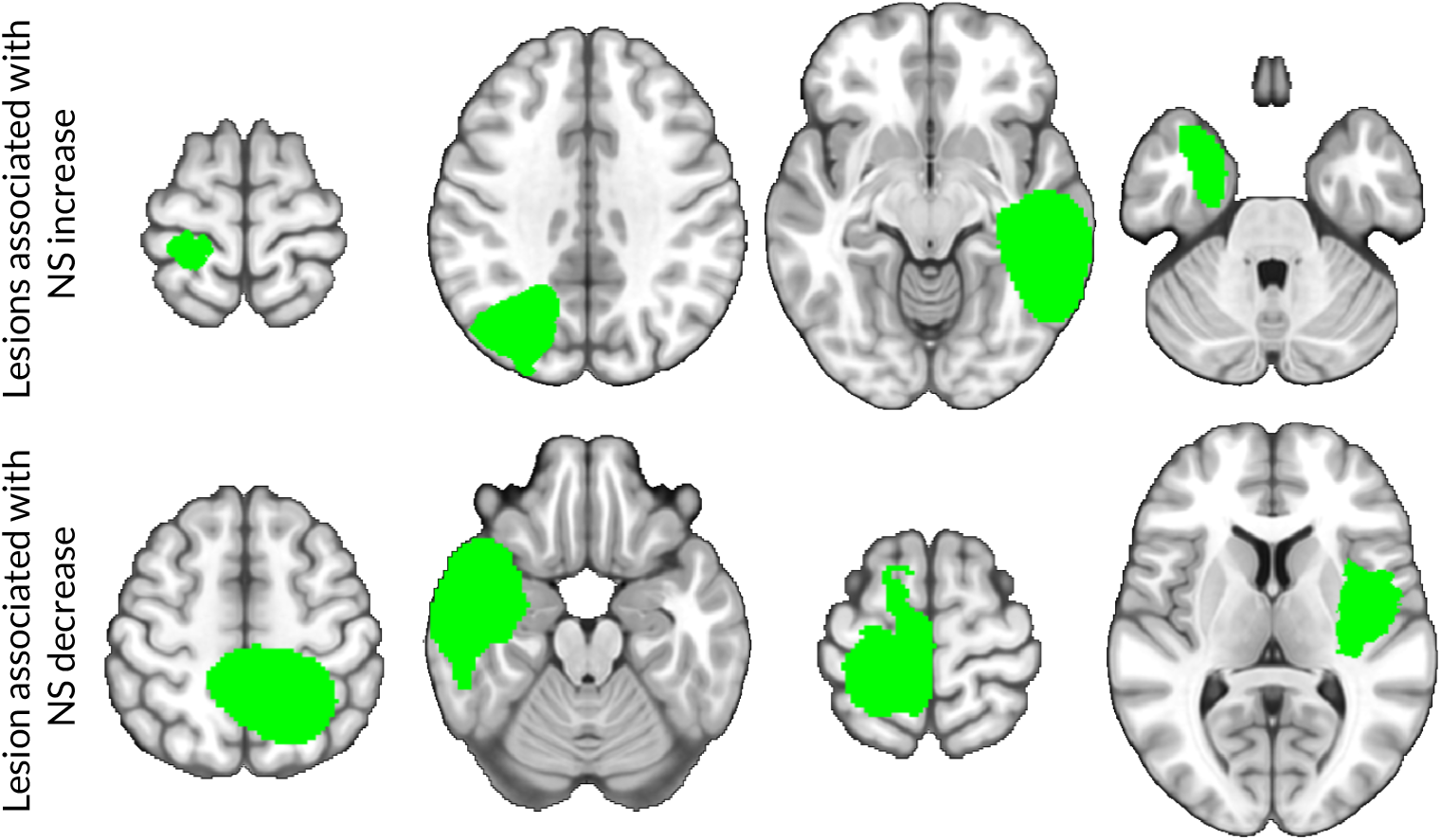
Example surgical lesions from two independent datasets associated with prominent increases and decreases in novelty seeking. Lesion locations were heterogeneous with respect to personality changes.

**Figure 2:**
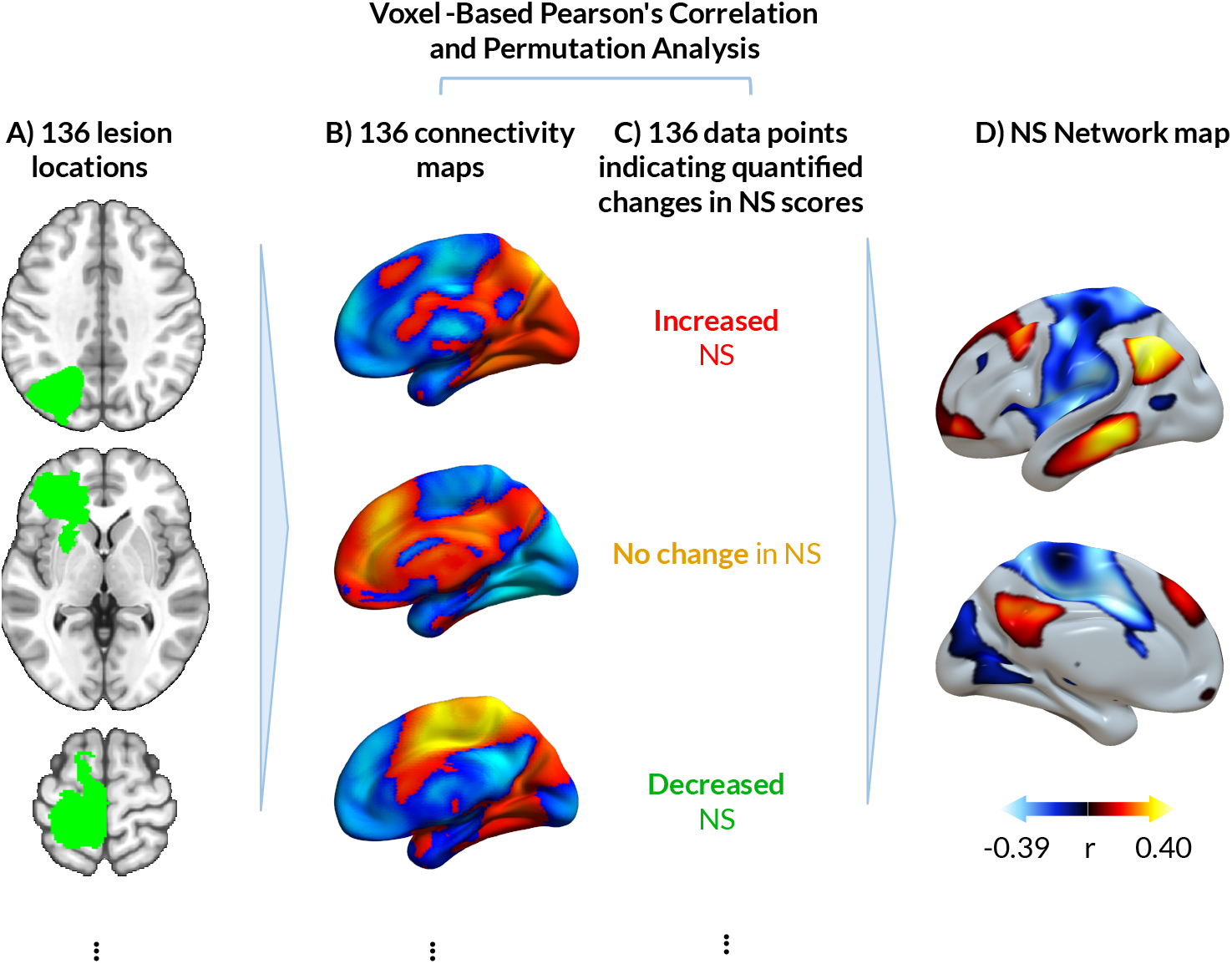
(a) Each lesion was normalized to a common MNI152 brain atlas. (b) Using a normative healthy human connectome (n=1000), a functional connectivity map was computed for each lesion seed. (c) Changes in novelty seeking following surgery were compared to each lesion’s connectivity to assess which connections are relevant to NS. (d) Changes in novelty seeking were more connected to a brain network including the cingulate and thalamus.

### Network mapping produced nearly identical maps from two independent datasets

Among all four TCI-defined temperament traits, only the lesion network mapping of NS produced highly identical network maps across both independent datasets (r = 0.80) (Fig. 3a). We computed statistical significance of this spatial correlation using permutation testing as in our prior work (Siddiqi et al., 2021). Briefly, we recomputed the spatial correlation after randomly permuting each patient’s neuroimaging with a different patient’s NS scores. We repeated this process 10,000 times and computed a p-value based on the percentage of cases in which the permuted spatial correlation was greater than the true value. Permutation testing confirmed that the two datasets yielded similar NS maps (p = 0.0061) (Figure 3a). This value survived Bonferroni correction across four temperament traits or across all seven personality traits (p < 0.05). We refer to the estimated lesion network map the “novelty seeking (NS) network” (Figure 2d). None of the other three temperament or three character traits showed significant cross-dataset replication.

**Figure 3:**
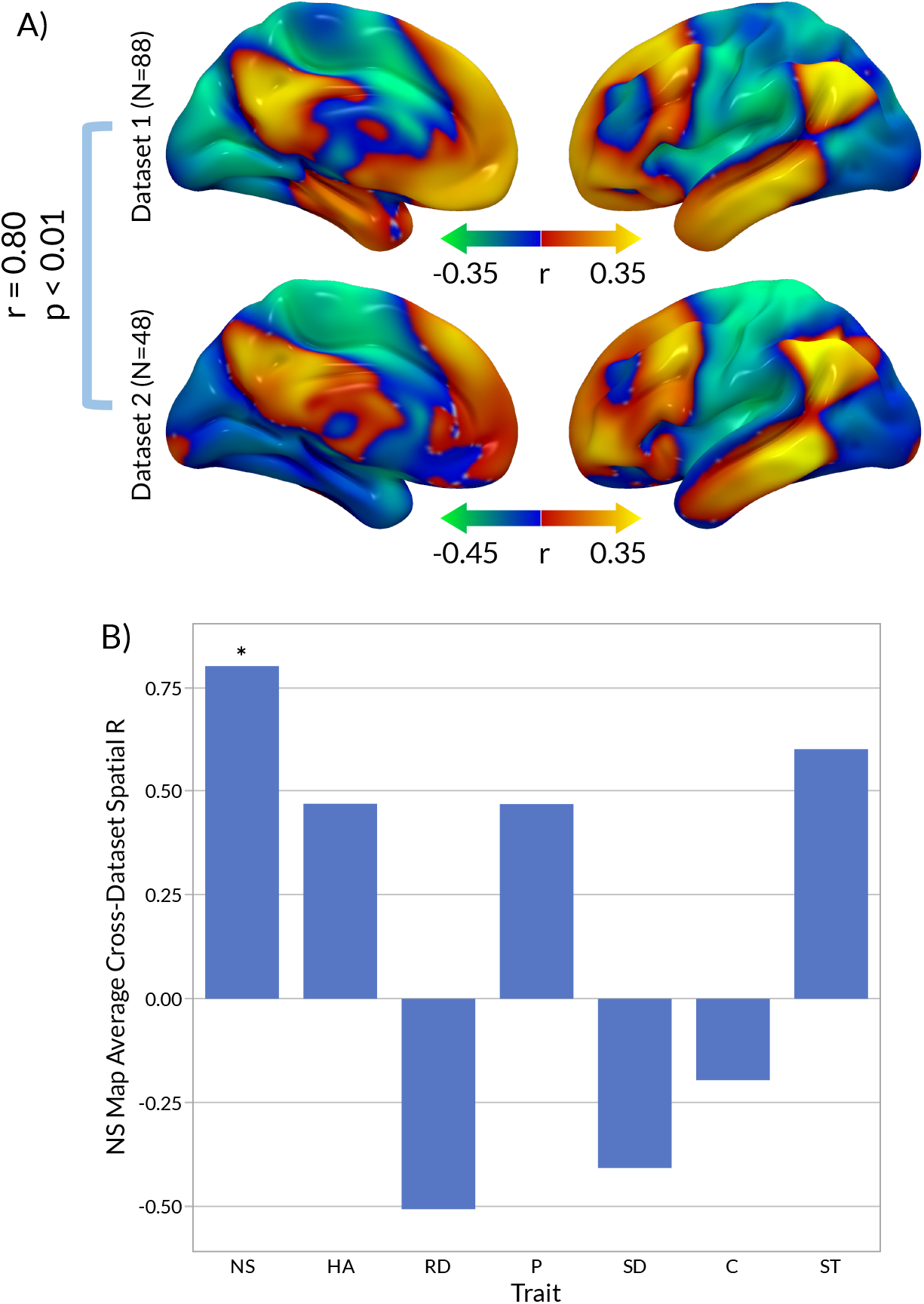
(a) The NS network maps generated from two independent datasets were highly similar (spatial r = 0.80). Permutation testing (10,000 permutations) indicated that this similarity was statistically significant (p = 0.0061). (b) A specificity analysis revealed that the NS maps were significantly more similar to each other than to all other TCI-based personality traits, on average (p = 0.012).

### Specificity to novelty seeking

To assess specificity, we recomputed the novelty seeking network after controlling for each of the remaining six TCI traits. The resulting NS specificity maps remained highly spatially correlated with each other across both datasets (r > 0.77, p < 0.012 across 10^5^ permutations).

We also assessed specificity by correlating the NS network from each dataset with control maps generated from the remaining TCI traits in the other dataset. The NS map from each dataset was more similar to the NS map from the other dataset than to the control maps from the other dataset. Permutation testing confirmed that the similarity between the NS maps across both datasets was significantly stronger than the mean similarity of each NS map with the control maps (p = 0.0117; Figure 3b).

### The predictive capacity of lesion connectivity maps for identifying change in NS

To further examine the predictive capacity of the identified NS brain network, we generated the NS network from each dataset individually, and used it to predict NS change in the other dataset. Lesion overlap with the NS network successfully predicted NS change in an independent sample (r = 0.2875, p < 0.001) (Fig. 4a). As a separate leave-one-out cross-validation, we generated an NS network based on all lesions across both datasets except for one. The excluded lesion connectivity map was spatially correlated with the leave-one-out overall network map. This process was repeated for each lesion, and the resulting connectivity estimates were correlated with each patient’s change in NS score. Lesion connectivity to the NS network was again predictive of change in novelty seeking (r = 0.2875, p < 0.0005) (Figure 4b). All of these analyses were controlled for lesion size and baseline novelty seeking score.

**Figure 4:**
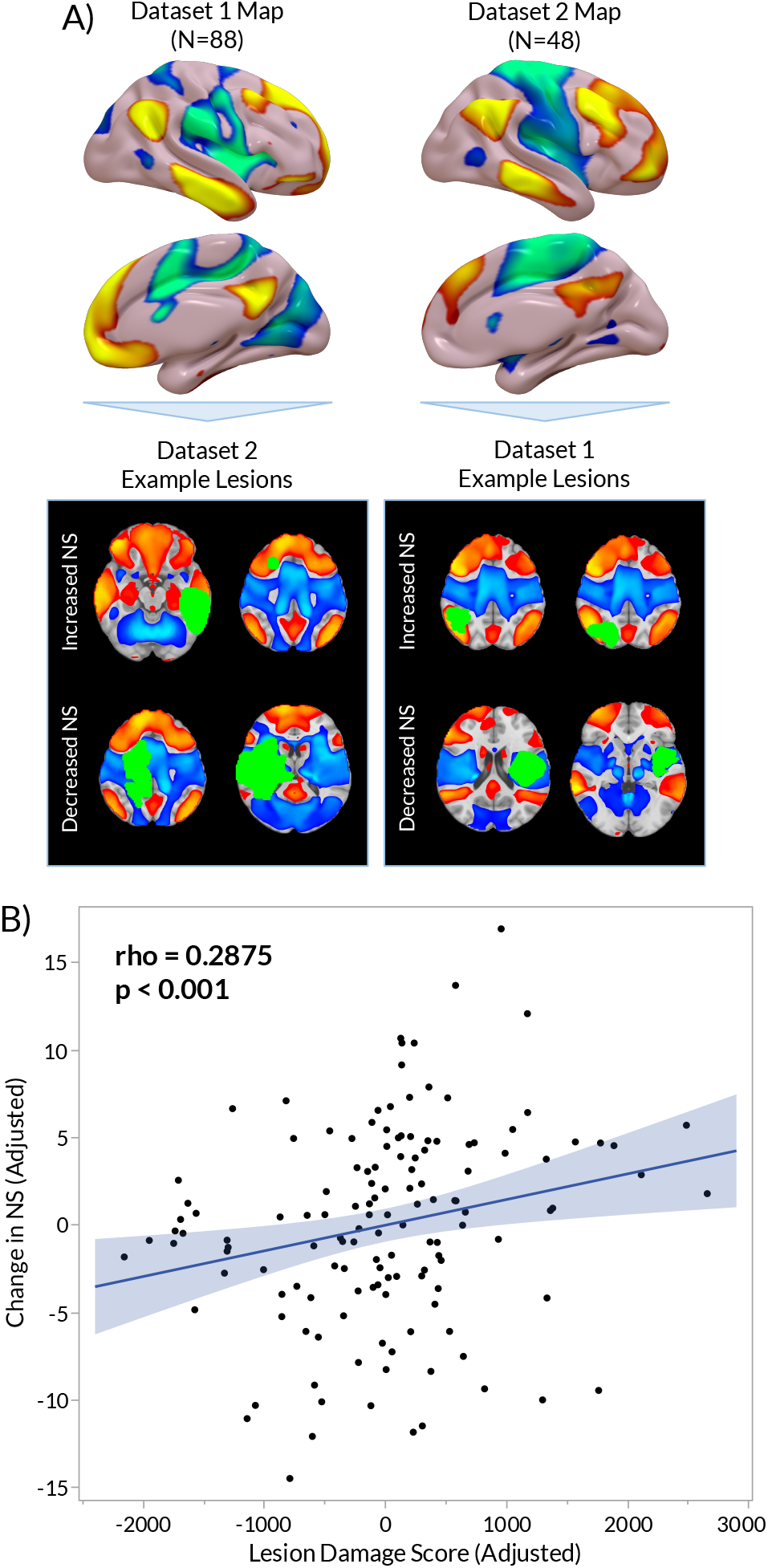
(a) In a dataset cross-validation, the lesion networks for each lesion in dataset 1 were compared to the network overlap map generated from dataset 2, and vice versa. (b) The degree of network overlap by each lesion strongly predicted change in trait novelty seeking (r = 0.29, p < 0.0005).

### Lesion connectivity to fMRI findings of novelty predict Novelty Seeking

To assess for generalizability beyond lesions, we used the Neurosynth platform (Yarkoni et al., 2011) to automatically meta-analyze neuroimaging studies related to the term “novelty”. This produced a map of voxels associated with significant activations compared to noise. We treated this map as an *a priori* seed. The connectivity of lesions to the meta-analytic novelty map was significantly predicted changes in NS (r = -0.2511, p < 0.005), again controlling for lesion size and baseline novelty seeking score.

### The NS map remains nearly unchanged when controlling for sex

To assess whether gender may have a significant impact on the NS network map, we repeated the lesion network mapping procedure using sex-specific connectome databases. Each lesion was sorted based on the subject’s sex and its connectivity was estimated using either a male-specific connectome (N = 500) or a female-specific connectome (N = 500), respectively. This yielded nearly identical network maps between datasets (spatial r > 0.97).

### Neuroanatomical specificity

To assess for neuroanatomical peaks in the NS network after multiple comparisons correction, we utilized Permutation Analysis of Linear Models (PALM) (Winkler et al., 2014) to construct voxel-wise general linear models, comparing unthresholded lesion connectivity maps to changes in NS score, with pre-lesion NS as a covariate. These models were used to identify regions associated with NS within an MNI brain mask. Significant clusters in this map were identified using threshold-free cluster enhancement (TFCE; Smith & Nichols, 2009) with a rigorous voxel-wise family-wise error (FWE) correction approach, set at p_FWE_ < 0.05 (two-tailed). Ten clusters consisting of greater than 10 voxels are described in Table 1. This analysis yielded a significant network of regions connected to lesions that modify NS (Figure 5), highlighting an extensive region in the dorsal cingulate.

**Figure 5:**
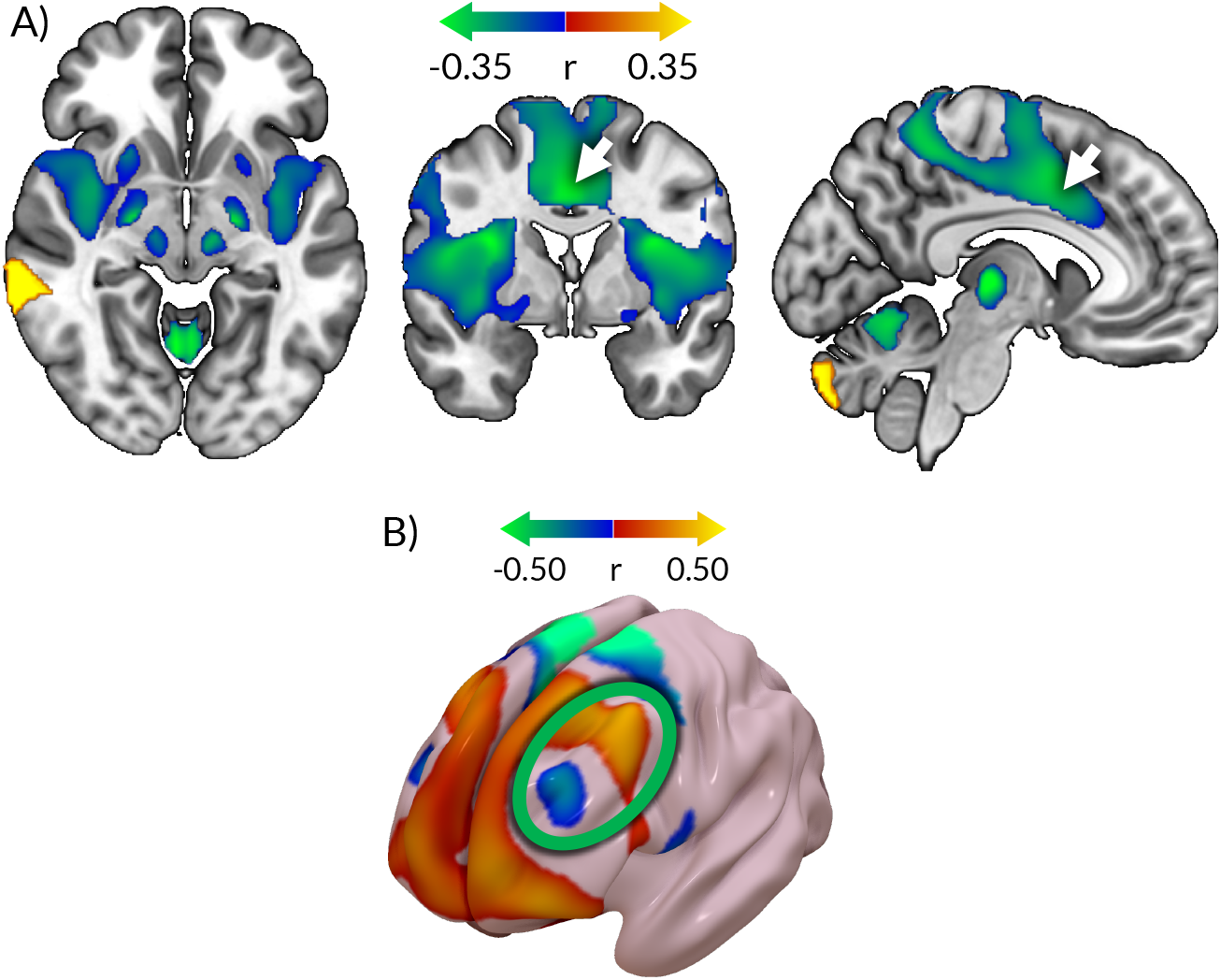
(a) The NS network map includes areas that survive a rigorous family-wise error (FWE) corrected threshold-free cluster enhancement (TFCE). The largest cluster in this map was a negative association in the dorsal cingulate. (b) The map resulting from a voxel-wise comparison of the FWE-corrected TFCE NS network map with the pre-computed connectome highlights which voxels share connectivity profiles with the NS network. This thresholded map reveals areas that may be targeted for brain stimulation to most directly activate or inhibit the network. The left dorsolateral prefrontal cortex, the most common clinical TMS target, includes both positive and negative areas on this map.

We also computed a more expansive network map using PALM’s two-tailed FWE-corrected threshold-free cluster enhancement, set at p_FWE_ < 0.05. TFCE yielded a significant network of regions connected to lesions that modify NS (Figure 5a), highlighting an extensive region in the dorsal cingulate.

### Potential targets for non-invasive brain stimulation

Using a precomputed whole-brain connectome database, we spatially correlated the TFCE FWE-corrected NS network map with each connectome voxel’s connectivity map. By definition, this analysis yields the similarity of each voxel’s connectivity profile to our NS network, yielding a targeting atlas of potential brain stimulation targets that would be expected to increase or decrease NS. The thresholded pre-computed NS connectivity map is shown in Figure 5b. Typical clinical TMS targets in the dorsolateral prefrontal cortex fell into both positive and negative areas within this map.

## Discussion

Using lesion network mapping, we have demonstrated that trait novelty seeking (as defined by the TCI) maps to a reproducible network. This Novelty Seeking network is specific relative to other personality traits. This lesion-derived NS map is not only causal (Fox, 2018), but also overlaps with prior neuroimaging findings related to the term “novelty”. Typical TMS targets for major depression overlapped with both the positive and negative portions of this map, suggesting that clinical TMS may have differential target-dependent effects on NS.

To our knowledge, this is the first study to demonstrate that a stable heritable personality trait maps to a reproducible brain network derived using causal sources of information. This network was centered on a large negative cluster in the dorsal anterior cingulate cortex, which has previously been associated with novelty seeking in correlative neuroimaging studies (S. Li et al., 2017). This region is generally considered to involved in sustained attention to task performance, and is part of a network that is variably called the cingulo-opercular, ventral attention, or salience network (Thomas Yeo et al., 2011). The negative association of novelty seeking with sustained attention is consistent with the established role of impulsivity as a core symptom in attention deficit-hyperactivity disorder (American Psychiatric Association, 2013).

Excitatory stimulation of positive regions in our network may be expected to decrease novelty seeking. This is consistent with multiple early-stage studies implicating the medial prefrontal cortex as a potential target for impulsivity (Kim & Lee, 2011; B. Li et al., 2020), which is strongly associated with novelty seeking. By contrast, excitatory stimulation of negative regions in our network may be expected to increase novelty seeking. This is consistent with a recent report showing that TMS to the DLPFC (at a site that is connected to the dorsal cingulate) led to increase in hypnotizability (Faerman et al., 2021), another stable personality trait that is associated with NS in some populations (Lichtenberg et al., 2004).

Strengths of our study include robustness across two independent datasets, rigorous cross-validation with rigorous permutation-based statistical assessments, specificity to NS versus other personality traits defined by TCI, and ability to control for pre-lesion scores, which is uncommon in lesion studies. An important limitation is the use of a self-report personality measure, which may not be the most precise metric of novelty seeking. The dataset was also limited to glioma resections, and resection of glial tissue may not have as direct of a behavioral impact as resection of neuronal tissue. Both of these limitations may introduce noise into the analysis and limit the observed effect size, so future studies may seek to use task-based novelty seeking measures along with different lesion types. Furthermore, while we propose a potential targeting atlas for TMS-induced changes in novelty seeking, this atlas should be considered to be a hypothesis rather than a conclusion, as it was not tested on TMS data. Future studies should seek to determine whether stimulation of different parts of this network can selectively induce differential changes in NS or related traits.

In conclusion, we derived and replicated a brain network that is functionally connected to lesions that can causally modify novelty seeking, a stable personality trait. Typical clinical TMS sites appear to be overlap with both positive and negative components of this network, suggesting that patients receiving routine clinical TMS may be experiencing target-dependent changes in novelty seeking. Future studies may employ these targets to augment personality disorders associated with excessive novelty seeking (such as borderline personality disorder) or inadequate novelty seeking (such as obsessive-compulsive personality disorder).

## Methods

### Primary dataset

Data from two distinct neurosurgical lesioning datasets were analyzed (Campanella et al., 2014; Urgesi et al., 2010). The first dataset (Dataset 1) comprised 88 participants (mean age: 47.4 ± 11.8 years), whereas the second dataset (Dataset 2) included 48 subjects (mean age: 48.1 ± 15.3 years).

Due to the potential for tissue regeneration post-surgery, lesion volumes were identified based on pre-surgery MRI scans. Lesion identification was performed by hand using the MRIcron and MRIcro software tools and normalized to the Montreal Neurological Institute (MNI) space (Campanella et al., 2014; Urgesi et al., 2010). All subjects underwent pre- and post-surgery examination of the Temperament and Character Inventory (TCI), a 240-item questionnaire for assessing personality traits. This inventory identifies seven personality subscales including Novelty Seeking, Harm Avoidance, Reward Dependence, Persistence, Self-Directedness, Cooperativeness, and Self-Transcendence.

### Outcome measures

The present study examined the impact of lesions on the seven personality sub-scales of the TCI by comparing pre- and post-operative measurements. To ensure the accuracy of the assessments, raw TCI scores were expressed as T scores to control for the potential influence of age and sex (Campanella et al., 2014; Urgesi et al., 2010).

### Lesion symptom mapping

Custom-built scripts in Python were utilized to estimate the whole-brain connectivity maps of the identified lesioned areas; a few example lesions are depicted in Figure 1. The normative human connectome database consisting of resting state scans from 1000 individuals was employed as a reference (Siddiqi et al., 2021). Time courses for voxels within each lesioned area were averaged for each normative connectivity map, and these averages were then correlated with the time course of all other voxels in the brain. Next, the resulting connectivity maps were averaged across all scans for each lesion to obtain a single functional connectivity map. Using Pearson’s partial correlation in MATLAB 2022b software package (MathWorks, Inc., Natick, Massachusetts, United States), the correlation between the change in personality scores and the connectivity maps of each lesion was estimated, while controlling for the effect of lesion volume and pre-surgery personality scores across all subjects. This generated a whole-brain network map for each personality trait, representing brain networks responsible for the alteration of that trait, where positive connections were associated with increases in that trait, and negative values demonstrated connections corresponding to decreases in the trait.

For each TCI-defined personality trait, network r-maps were also generated individually for each of the two independent datasets, using the procedure described above. Each map was then Fisher z-transformed, using the MATLAB atanh function, and spatially correlated across the two datasets using a Pearson’s correlation to determine the strength of the association. Significance of the association was assessed using a permutation testing approach. For each permutation, each dataset’s trait network map was regenerated by randomly shuffling the order of each lesion’s connectivity map relative to the change in NS score and covariates. The permuted network maps were then Fisher z-transformed and spatially correlated, and the resulting r-values were compared to the true spatial correlation between the datasets. This process was repeated for 10,000 permutations, allowing for a permuted p-value to be generated based on the number of permuted r-values that were greater than the true r-value. The personality trait network map most significantly reproducible across Datasets 1 and 2 was selected for further examinations in the rest of this study.

### Specificity relative to other personality traits

To determine whether the estimated NS map is specific relative to other personality traits, the network map was also controlled for changes in each of the other TCI-defined personality traits. To this end, a network map of novelty seeking was produced within each dataset by correlating each lesion’s connectivity map with change in NS scores in that dataset, while controlling for pre-lesion NS scores and lesion size. For specificity analysis, each map was additionally controlled for changes in all other TCI-defined personality traits. Following a similar procedure, we created 12 distinct maps for each of the TCI-based personality traits. The respective maps for Datasets 1 and 2 were Fisher z-transformed and spatially correlated to determine whether the cross-dataset similarity may be due to any of the other personality traits. Additionally, a permutation testing approach was employed as described above. This produced a p-value for the true spatial correlation of the NS maps after controlling for the effect of each of the remaining traits.

Specificity was also assessed by comparing the similarity of Dataset 1 and 2 NS maps versus the maps of each remaining TCI personality trait. The Fisher-transformed Dataset 1 NS map was spatially correlated with each of the six other Fisher-transformed trait network maps of Dataset 2 and vice versa for the Dataset 2 NS map. The differences between the spatial correlation of the NS maps and those of the NS maps compared to all other maps were averaged. To test the significance of this average difference, permutation testing was employed as above. The proportion of times the permuted average difference exceeded the true average difference was calculated, producing a p-value for this analysis. Additionally, this spatial correlation difference and permutation testing process was repeated for each trait individually to determine against which traits the NS map was more significantly specific than expected by chance.

### Dataset cross-validation

To assess the predictive capacity of the NS lesion network map in predicting changes in NS, a dataset cross-validation pipeline was employed. Each lesion from Dataset 1 was used to individually mask the Fisher z-transformed lesion network map produced from Dataset 2. The sum of the r-values for voxels confided in the masked area of the map provided a single network damage score for each lesion, which was treated as a predictor. The same procedure was repeated for the lesions of Dataset 2, which were used to mask the network map produced from Dataset 1. The correlation between the resulting network damage scores and each lesion’s change in NS scores was estimated, controlling for the effect of pre-lesion NS score. Lesion volume was not included as a covariate as volume of damage may be part of the mechanism by which a lesion has an impact on the network.

As a secondary assessment, we utilized a leave-one-out analysis where all lesion connectivity maps, except for one, were combined, and a new leave-one-out network map was estimated based on the remaining connectivity maps using the procedure described above. The excluded map was then correlated with the resulting Fisher z-transformed NS network r-map to produce a predictor. This procedure was repeated for each lesion. Controlling for the effect of pre-lesion NS score and lesion size, the resulting Fisher z-transformed predictors were correlated with each lesion’s change in NS and the strength and significance of the relationship was evaluated.

### Generalizability beyond brain lesions

Generalizability of the NS network map beyond brain lesions was assessed using Neurosynth, a large-scale tool for automated meta-analysis of neuroimaging data associated with specific terms (Yarkoni et al., 2011). Neurosynth was used to extract activation coordinates from studies related to the term “novelty”. This produced a map indicating which voxels are consistently activated across relevant studies at a rate differing from a uniform grey matter distribution. The lesion connectivity maps were spatially correlated with each lesion’s connectivity map to produce a predictor. This predictor was then Fisher-transformed and correlated with each lesion’s change in NS, controlled for the effect of pre-lesion score.

### Controlling for sex

The impact of sex on the NS network map was assessed by recomputing the lesion connectivity maps for both datasets using sex-specific connectomes. Two normative human connectomes consisting of resting state scans from 500 male and female individuals each were used. Lesions were separated based on the subject’s sex and the lesion network mapping approach was employed as described above using the respective sex-specific connectome. The resulting sex-specific lesion connectivity t-maps were once again spatially correlated with change in NS, controlling for pre-lesion NS score and lesion volume, to produce a sex-adjusted NS network map for each dataset. The original Fisher z-transformed Dataset 1 NS map was then spatially correlated with the Fisher z-transformed sex-adjusted map for Dataset 1 to determine the impact of adjusting for sex in the development of the lesion network map. This was repeated for Dataset 2 and the combined dataset maps.

### Localization of significant clusters

Lesion connectivity maps causing changes to NS were compared using a general linear model and permutation testing to assess the significance of each voxel in the NS network (Permutation Analysis and Linear Modeling software included in FSL 3.2.0; Winkler et al., 2014). The resulting t-map was Fisher z-transformed and processed using threshold-free cluster enhancement. TFCE is a method of identifying significant regions by utilizing information regarding the spatial extent of significant voxels without having to rely on cluster-level thresholds, thus avoiding false positive results, as previously reported (Eklund et al., 2016; Smith & Nichols, 2009). Family-wise error (FWE) correction was applied to correct for multiple comparisons, and the corrected p-map was thresholded at p < 0.05. Using the cluster command included in FSL 6.0.4, significant clusters larger than 10 voxels were isolated and are listed in Supplemental Table 1. Relevant brain regions were identified using the Talairach Daemon Labels for the cerebrum and FNIRT Cerebellar Atlas for the cerebellum, which are included as standard atlases in the FSLeyes (version 1.3.0) image visualization software.

### Identification of Sites for Stimulation

A previously generated pre-computed connectome was used to highlight relevant targets for non-invasive stimulation of the NS network. This version of the connectome was derived from the normative connectome of 1000 healthy controls, consisting of maps of every voxel’s connectivity to every other voxel in the brain. Using custom-built Python scripts, the TFCE FWE-corrected masked NS network map was spatially correlated with each voxel’s connectivity profile. Each correlation value was assigned to its respective voxel to produce a new map highlighting which brain voxels share connectivity profiles most similar to that of the NS network. In turn, this highlights which brain areas may be best targeted with stimulation to activate an NS network-like network. This pre-computed connectome-derived map was masked by the prefrontal cortex and presented in surface space to spotlight those areas most easily targeted by interventions such as TMS.

