## Supplemental Table 1 for "Lesion network localization of a stable personality trait"

| Supplemental Table 1. Significant clusters returned in the TFCE FWE-corrected NS network map | | | |
| --- | --- | --- | --- |
| Significant Clusters | | | |
| Brain Regions | Significance (FWE-corrected) | Volume (mm^3^) | Peak Voxel Coordinates (MNI) |
| Dorsal cingulate, thalamus, and globus pallidus - bilateral | p < 0.005 | 256,080 | X: 57, Y: 50, Z:46  (cingulate: X: 45, Y: 66, Z:55) |
| Cerebellum (culmen, declive) | p < 0.05 | 12,504 | X: 46, Y: 30, Z: 30 |
| Angular gyrus - right | p < 0.05 | 8,488 | X: 65, Y: 35, Z: 53 |
| Cerebellum (crus II) - left | p < 0.05 | 6,200 | X: 38, Y: 22, Z: 12 |
| Middle temporal gyrus - right | p < 0.05 | 5,768 | X: 80, Y: 42, Z: 32 |
